# Mitigating inflammatory bone loss in post-menopausal osteoporosis via targeting the IL-9 producing osteoclastogenic Th9 cells

**DOI:** 10.1101/2023.11.25.568680

**Authors:** Leena Sapra, Chaman Saini, Pradyumna K. Mishra, Naibedya Chattopadhya, Rupesh K. Srivastava

## Abstract

Recent discoveries have established the pivotal role of IL-9-secreting Th9 cells in a wide spectrum of inflammatory and autoimmune diseases. However, little is known about how Th9 cells contribute to the etiology of inflammatory bone loss in post-menopausal osteoporosis (PMO). We observed that IL-9 has a pathological impact on inflammatory bone loss in ovariectomized (Ovx) mice. Our in vivo temporal kinetics analysis further revealed that estrogen deprivation increased the release of IL-9 from Th which in turn enhances the IL-17-producing Th17 cells. Both ex vivo and in vivo studies corroborated these findings in Ovx mice, as estrogen diminishes IL-9’s effect on the differentiation of Th17 cells as well as the potential of Th9 cells to produce IL-9. Mechanistically, Th9 cells in an IL-9-dependent manner enhance osteoclastogenesis and thereby establish themselves as a novel and independent osteoclastogenic Th subset. Blocking IL-9 improves bone health in Ovx mice by inhibiting the differentiation and function of both osteoclasts and Th9/Th17 cells. Our clinical findings further attested to the osteoporotic role of Th9 cells in post-menopausal osteoporotic human subjects. Collectively, our study establishes IL-9-secreting Th cells as the critical regulator of bone loss observed in PMO and highlights the fundamental implications of IL-9/Th9 targeted immunotherapies as an innovative approach for the treatment of inflammatory bone loss observed in osteoporosis.

## Introduction

Osteoporosis is a systemic skeletal illness that is principally distinguished by a loss of bone mechanical strength (BMS) and bone mineral density (BMD), thereby raising the risk of fragility-related fractures in the wrist, hip, and spine ^1,2^. It is the 4^th^ most burdensome chronic disease after ischemic heart disease, dementia, and lung cancer, impacting over 500 million people globally ^3^. Postmenopausal osteoporosis (PMO) is a common skeletal disorder that increases the risk of fragility fractures and attendant morbidity. Postmenopausal bone loss is linked to chronically low-grade inflammation primarily because of the role played by estrogen on the immune system ^4^. Estrogen has anti-inflammatory effects and regulate the activity of immune cells in manners that downregulates the production of pro-inflammatory cytokines such as IL-6, TNFα, and IL-17 ^5^. These pro-inflammatory cytokines activate osteoclasts leading to greater bone breakdown over formation.

Moreover, the bone loss caused by low estrogen and PTH has been linked to IL-17 producing Th17 cells ^6,7^, as enhanced expression of IL-17 stimulates the expression and activity of RANKL, resulting in the enhanced bone loss ^7^. We and others have demonstrated that the disruption or breakdown of the homeostatic balance between the osteoclastogenic Th17 cells and anti-osteoclastogenic Tregs cells is one of the major underlying mechanisms of bone loss in osteoporosis ^8–10^. The disruption of Th17-Treg balance is also responsible for the inflammatory bone loss caused by psoriasis, rheumatoid arthritis (RA), periodontal disease, and inflammatory bowel disease (IBD) ^11–13^. Th cells develop into functionally varied subsets, including Th1, Th2, Th17, Th22, follicular T helper cells (Tfh), and Tregs. Distinct subsets of Th cells, each with their own unique cytokine profile and functional properties, define the nature of their immune responses. One such subset of T helper cells that mainly secretes the pro-inflammatory IL-9 cytokine was discovered and was later named as “Th9 cells” in 2008 ^14^. Th9 cell differentiation is predominantly induced by the cytokines transforming growth factor-beta 1 (TGFβ-1) and IL-4. Additionally, forkhead family transcription factor (FOXO1) and interferon regulatory factor 4 (IRF4) are essential for the growth and function of Th9 cells ^15–17^

Th9 cells enhance immunological tolerance and afford protection against parasitic infections^18^. They also have potent anti-tumor immunity, making them a potential candidate for cancer immunotherapy. Conversely, Th9 cells contribute to widespread allergic inflammation, asthma, and autoimmune diseases, highlighting their pathogenic roles in a range of immunological disorders ^19^. Furthermore, the presence of Th9 cells and the production of IL-9 in the synovial fluid and PBMCs of RA patients indicate that IL-9 may be involved in the inflammatory processes within the joints affected by RA ^20^. These findings demonstrate that inhibiting IL-9 signaling may be a viable strategy for limiting the inflammatory deterioration of bone, not just in RA but also in PMO. However, the regulation of Th9-IL-9 system in PMO is largely unknown.

We hypothesized that estrogen deficiency after menopause will lead to dysregulation of Th9 cells and IL-9 production, resulting in increased bone resorption. We addressed this hypothesis by examining the Th9-IL-9 system in both a mouse model of postmenopausal osteoporotic mice model (bilateral Ovx) and postmenopausal women with osteoporosis to better understand the connection between this immune pathway and the disease. Our data collectively demonstrate a distinct pathological role of Th9 cells in triggering inflammatory bone loss in estrogen deficient conditions, thereby opening novel therapeutic avenues for IL-9 targeted immunotherapy in post-menopausal osteoporosis.

## Results

### IL-9 levels negatively impact bone mass and microarchitecture

The study design is illustrated in **(Fig S1A)**. Bilateral Ovx in mice resulted in nearly 3-fold decline in circulating E2 levels compared with the ovary-intact (sham surgery) control mice **(Fig S1B)**. SEM, histological and μCT analysis of femoral bones revealed enhanced lacunae along with a significant decrease in BMD and BV/TV in the Ovx mice compared with the sham group suggesting reduced bone mass and volume **(Fig. S1C-H)**. Moreover, there was deterioration of trabecular microarchitecture in the Ovx group evident from reduced Tb. N, Tb. Th and increase in Tb.Sp **(Fig S1I-K) (Fig. S2A-D)**. This data suggests successful induction of osteoporotic mice model in Ovx mice. Serum levels of inflammatory cytokines such as IL-6, IL-17 and TNF-α were markedly enhanced while the levels of anti-inflammatory cytokines including IL-4, IL-10 and IFN-γ were decreased in the Ovx group compared with sham **(Fig. 1A-B)**, confirming the results of our earlier reports ^21,22^. Interestingly, serum IL-9 level was robustly increased in the Ovx mice compared with the sham and there was a significant negative correlation between IL-9 level and BMD **(Fig. 1C-D)**. In the bone marrow, the mRNA levels of IL-9 and IL-9R were significantly enhanced in the Ovx group compared with sham **(Fig. 1E-F)**. Moreover, Foxo-1, the transcription factor that supports that differentiation of naïve CD4^+^ cells to Th9 cells were also increased in the Ovx group **(Fig. 1G)**. These data suggest a positive feedback in bone marrow cells where IL-9 increase is accompanied by concomitant increase in its receptor and Foxo-1. From these data, it appears that estrogen deficiency promotes Th9 cell differentiation by increasing Foxo-1 expression. Th9 cells which produce IL-9, in turn further increases IL-9 and its receptor, likely amplifying Th9 cell differentiation in Ovx bone marrow.

**Fig 1:**
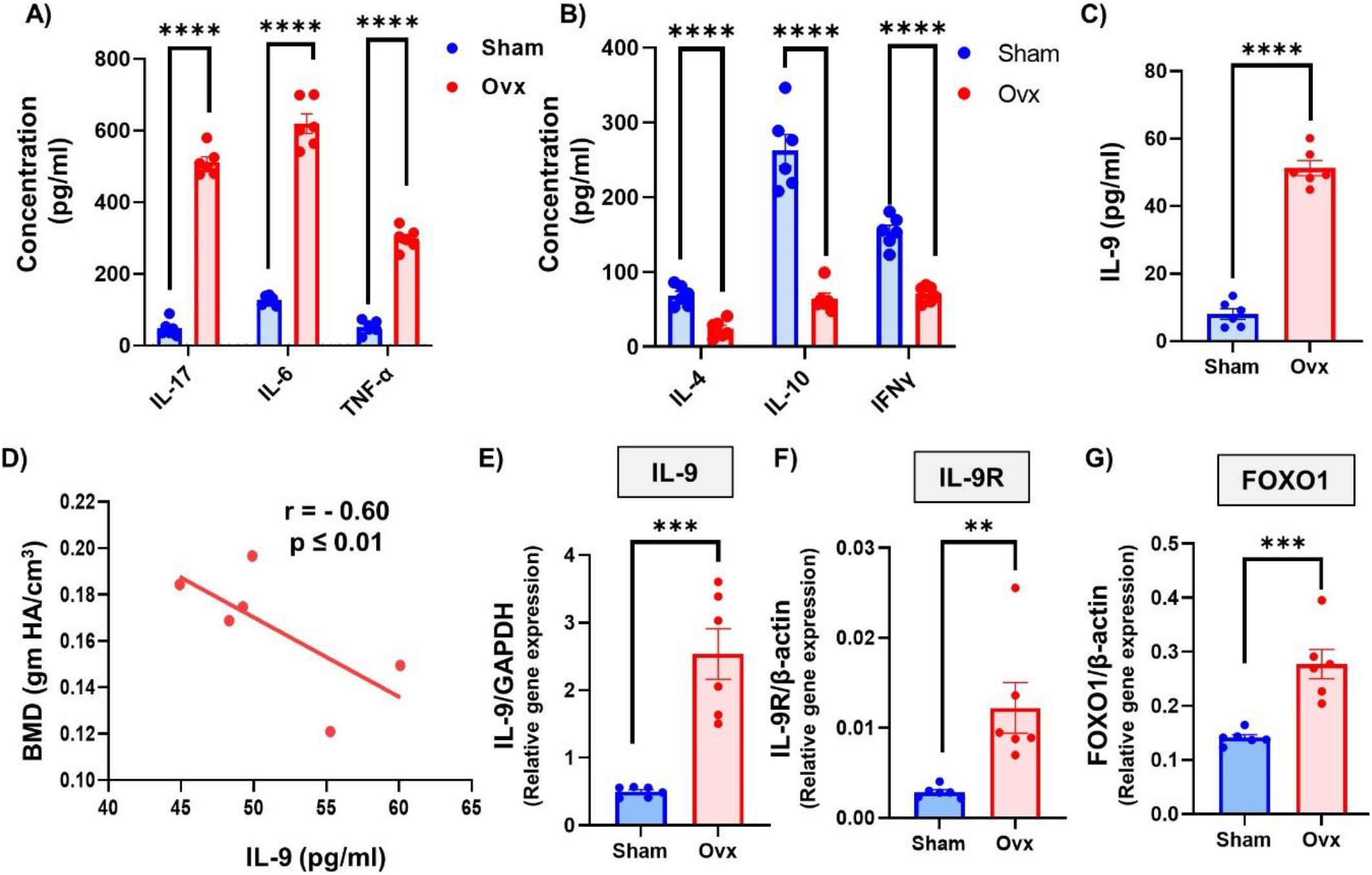
IL-9 cytokine levels negatively correlate with the bone mineral density (BMD): A) Levels of inflammatory cytokine in the sera of sham and Ovx. B) Levels of anti-inflammatory cytokines in the sera. C) Bar graphs representing IL-9 levels. D) Correlation between IL-9 cytokine and bone mineral density (BMD). E ) Relative gene expression of IL-9 gene. F) Relative gene expression of IL-9R. G) Relative gene expression of FOXO-1. For correlation, Pearson correlation coefficient was determined. Data are expressed as mean ± SEM. Data were analyzed by unpaired student t test and analyzed by one way ANOVA. *P ≤ 0.05, **P ≤ 0.01, ***P ≤ 0.001, ****p ≤ 0.0001) compared with the indicated group.

### Th9 and Th17 subsets are the major source of IL-9 cytokine

IL-9 is a pleiotropic cytokine, and is produced by various immune cells including ILC2, Th2, Tregs and Th17 alongside their specific cytokines. Next, we sought to pinpoint the specific cellular source of IL-9 via flow cytometry and found that ILC2, Th2, and Tregs were not significant sources of IL-9 in Ovx mice **(Fig 2A-F)**.

**Fig 2:**
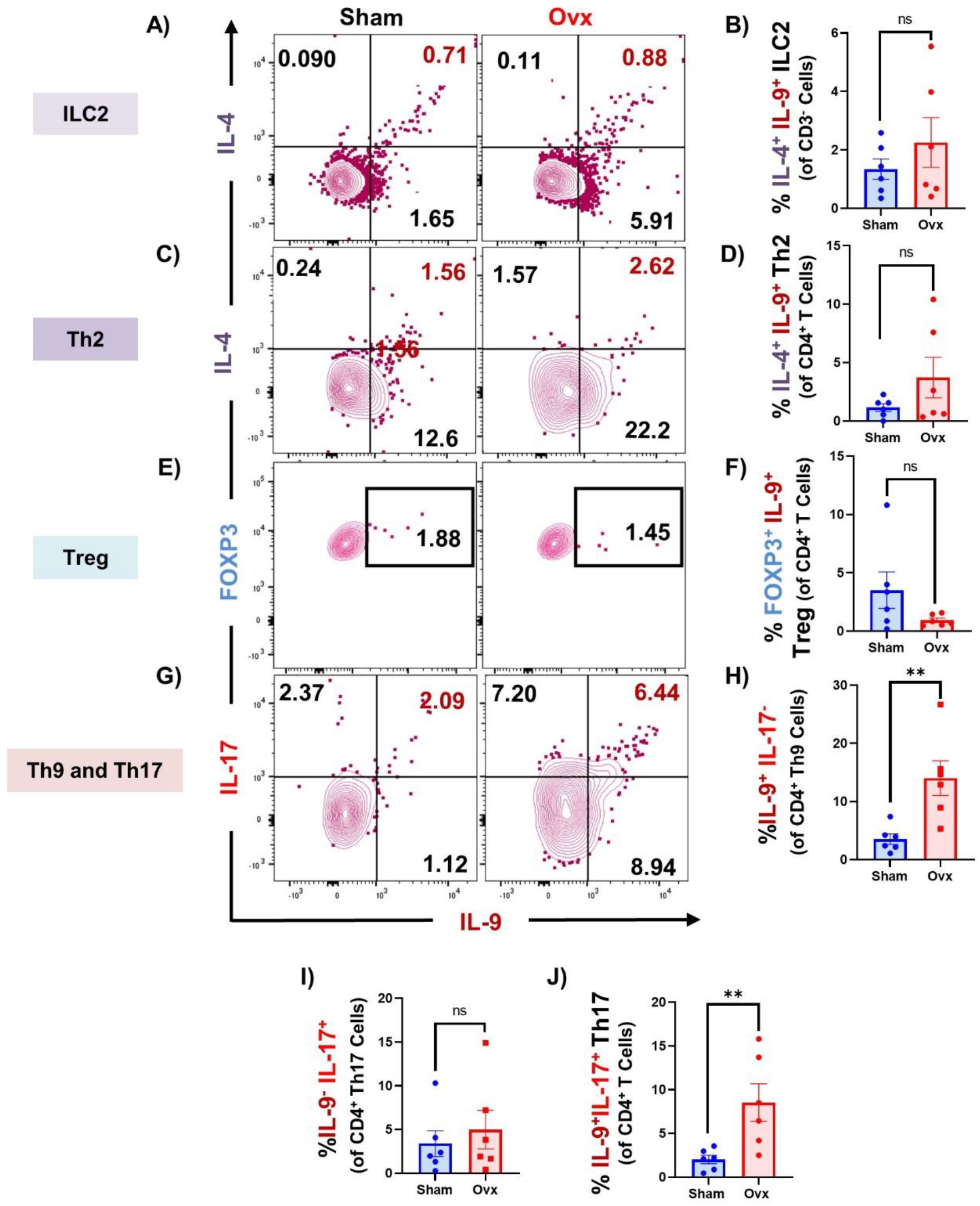
Th9 and Th17 subsets are the major source of IL-9 cytokine: A) Contour plot representing the percentage of CD3^-^IL-4^+^IL-9^+^ ILC2. B) Bar graphs representing the percentage of CD3^-^IL-4^+^IL-9^+^ ILC2. C) Contour plot representing the percentage of CD4^+^IL-4^+^IL-9^+^ Th2. D) Bar graphs representing the percentage of CD4^+^IL-4^+^IL-9^+^ Th2. E) Contour plot representing the percentage of CD4^+^Foxp3^+^IL-9^+^ Treg. F) Bar graphs representing the percentage of CD4^+^Foxp3^+^IL-9^+^ Treg. G) Contour plot representing percentage of CD4^+^IL-9^+^IL-17^+^ Th cells. H) Bar graphs representing percentage of CD4^+^IL-9^+^IL-17^+^ Th17 cells. I) Bar graphs representing percentage of CD4^+^IL-9^+^ Th9 cells. J) CD4^+^IL-IL-17^+^ Th17 cells. Data are expressed as mean ± SEM. Data were analyzed by unpaired student t test and analyzed by one way ANOVA. *P ≤ 0.05, **P ≤ 0.01, ***P ≤ 0.001, ****p ≤ 0.0001) compared with the indicated group.

Furthermore, the percentage of both IL-9^+^IL-17^-^ Th9 cells and IL-9^+^IL-17^+^ Th17 cells were significantly enhanced without any change in the percentage of IL-17^+^ Th17 cells in Ovx mice **(Fig. 2G-J)**. We and others have previously demonstrated that Th17 cells are pivotal in the pathogenesis of osteoporosis ^10^. Consistent with these findings, both *Il-17* mRNA levels and the percentage of IL-17 secreting Th17 cells were significantly enhanced in Ovx group over the sham group **(Fig. 3A-C)**. To further interrogate the likely contribution of IL-9^+^ Th cells in the differentiation of Th17 cells in PMO, we sought to perform kinetic studies at various time points (15, 30 and 45 days). Of note, we observed that IL-9 levels positively correlated with IL-17 **(Fig. 3D)**, and there was a time-dependent (15, 30 and 45 days) increase in Th9 and Th17 cells in Ovx mice **(Fig. 3E)**. In *ex vivo* experiments, the stimulatory effect of IL-9 in the differentiation of Th17 cells was completely abolished by E2 **(Fig. 3F-H)** which underscored the pathogenic link between estrogen deficiency and the activation of IL9-IL-17 axis thereby accelerating the onset and severity of bone loss in PMO.

**Fig 3:**
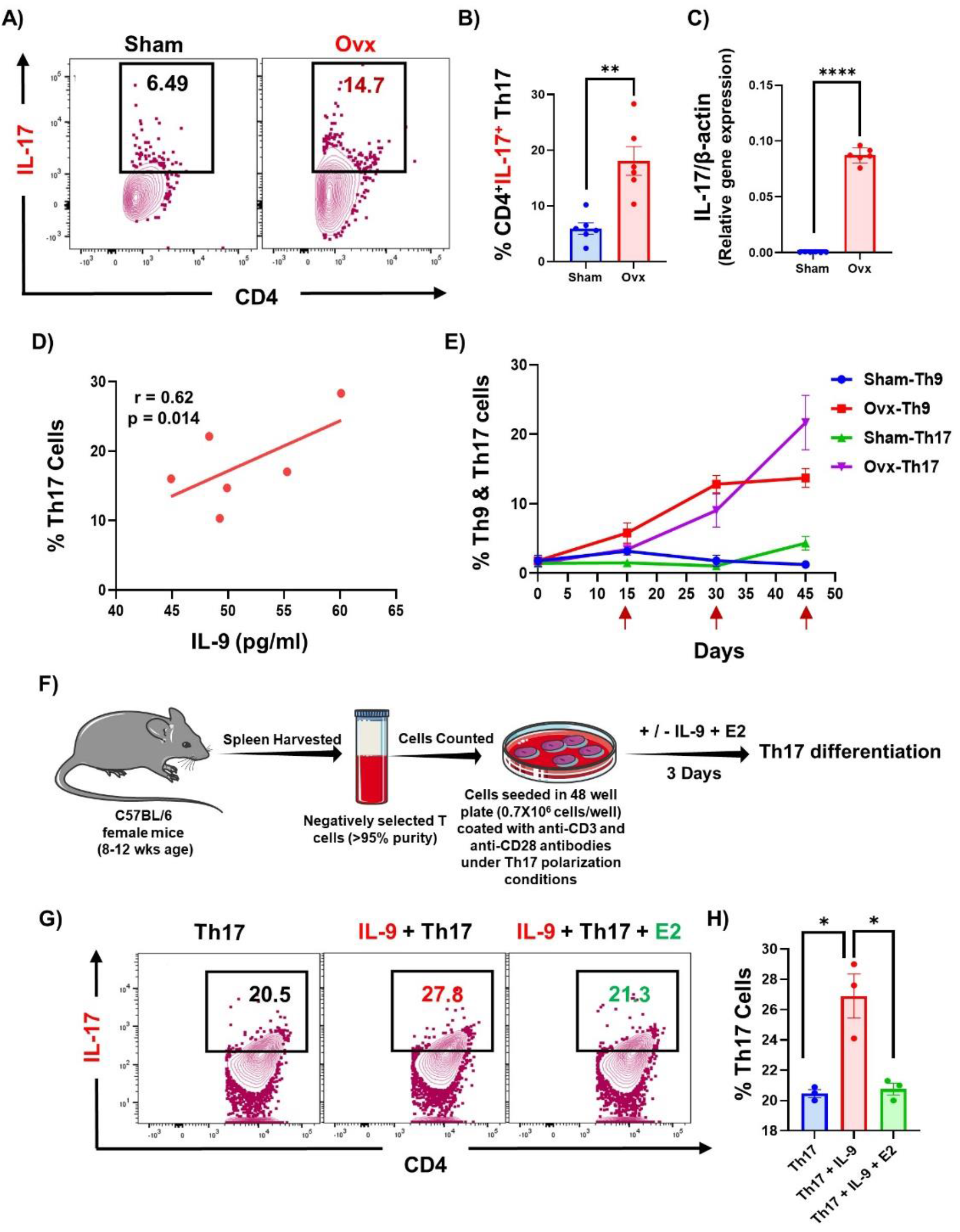
IL-9 enhance the differentiation of Th17 cells: A) Contour plot representing the percentage of CD4^+^IL-17^+^ Th17 cells. B) Bar graph representing the percentage of CD4^+^IL-17^+^ Th17 cells. C) Bar graphs showing relative expression of *Il-17a* gene. D) Correlation between percentage of CD4^+^IL-17^+^ Th17 cells and IL-17 gene. E) Line graphs percentage of CD4^+^IL-17^+^ Th17 cells and CD4^+^IL-9^+^ Th9 cells. F) Experimental layout showing Th17 cell differentiation. G) Contour plot representing CD4^+^IL-17^+^ Th17 cells in the presence or absence of IL-9 and E2. H) Bar graphs representing percentage of CD4^+^IL-17^+^ Th17 cells. Data are expressed as mean ± SEM. Data were analyzed by unpaired student t test and analyzed by one way ANOVA. *P ≤ 0.05, **P ≤ 0.01, ***P ≤ 0.001, ****p ≤ 0.0001) compared with the indicated group.

### Th9 cells are enhanced in post-menopausal osteoporotic conditions

We next studied the IL-9^+^ Th cells in BM and secondary lymphoid organs (spleen and mesenteric lymph nodes-MLNs). There was an increasing trend of IL-9^+^Th9 cells (both percentage and MFI) in the BM of Ovx mice **(Fig. 4A-E)** at day 15 post-Ovx, which was significantly enhanced at day 30 and day 45 **(Fig. 4F-L)**. Similar results were also observed in both spleen and MLNs at day 45 post-Ovx **(Fig. S3A-H and Fig. S4A-H)**. In *ex vivo* experiment, E2 dose-dependently inhibited the differentiation of Th9 cells **(Fig.5A-D)**. Thus, it appears that the presence of E2 keeps Th9 population in check, likely by reducing their proliferation and/or activation thereby modulating bone health.

**Fig 4:**
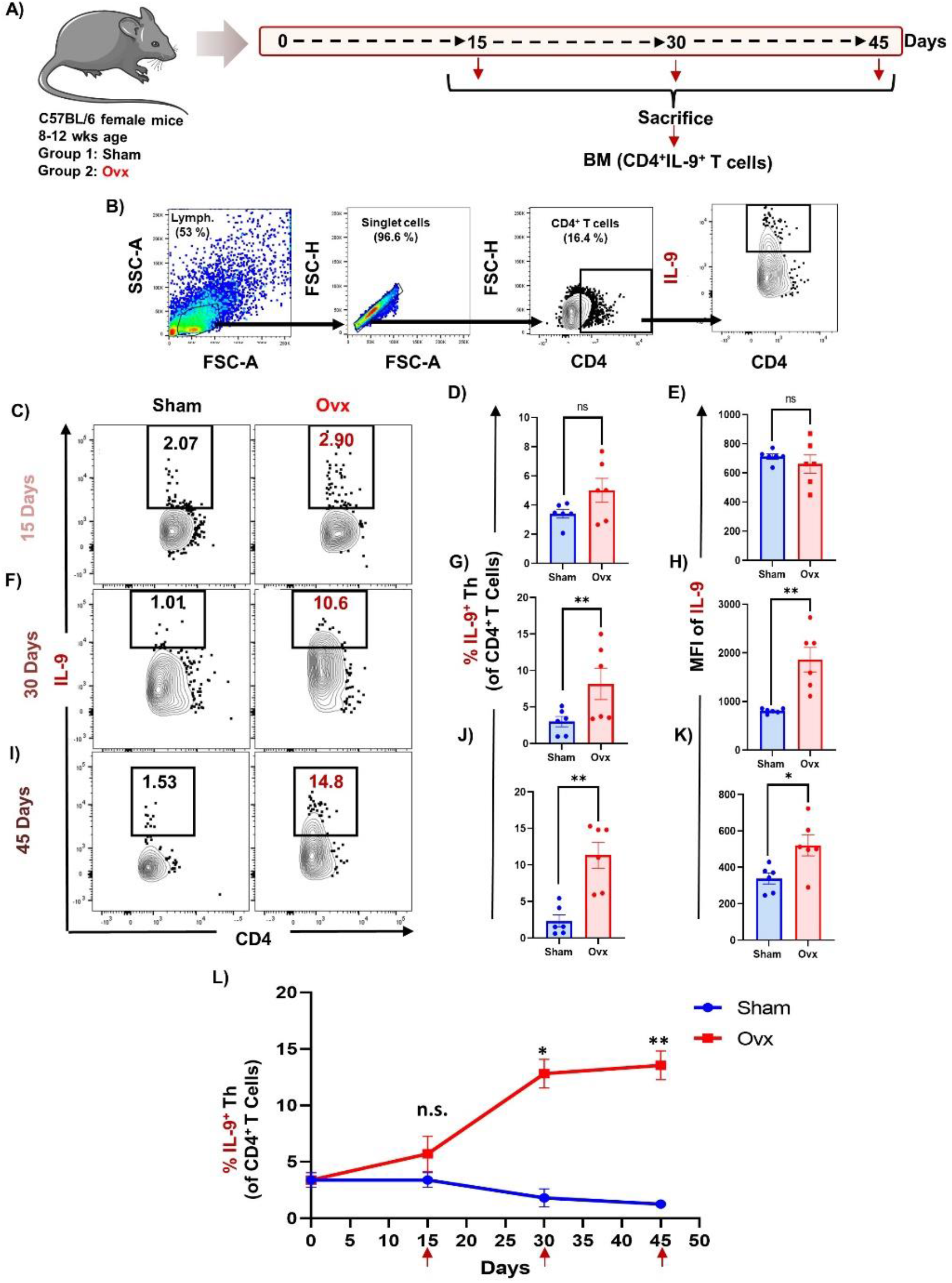
Th9 cells are enhanced in post-menopausal osteoporotic conditions: Cells from Bone Marrow (BM) of sham and ovx groups were harvested and analysed by flow cytometry for percentage of Th9. A) Gating strategy followed for data analysis. B) Contour plot depicting percentages of CD4^+^IL-9^+^ Th9 cells in BM at 15 days. C) Bar graph representing percentages CD4^+^IL-9^+^ Th9 cells in sham and ovx. D) Contour plot depicting percentages of CD4^+^IL-9^+^ Th9 cells in BM at 30 days. E) Bar graph representing percentages CD4^+^IL-9^+^ Th9 cells in sham and ovx. F) Contour plot depicting percentages of CD4^+^IL-9^+^ Th9 cells in BM at 45 days. G) Bar graph representing percentages CD4^+^IL-9^+^ Th9 cells in sham and ovx. H) Line graphs representing the percentage of CD4^+^IL-19^+^ Th9 cells at day 15, 30 and day 45. Data are expressed as mean ± SEM. Data were analyzed by unpaired student t test and analyzed by one way ANOVA. *P ≤ 0.05, **P ≤ 0.01, ***P ≤ 0.001, ****p ≤ 0.0001) compared with the indicated group.

**Fig 5:**
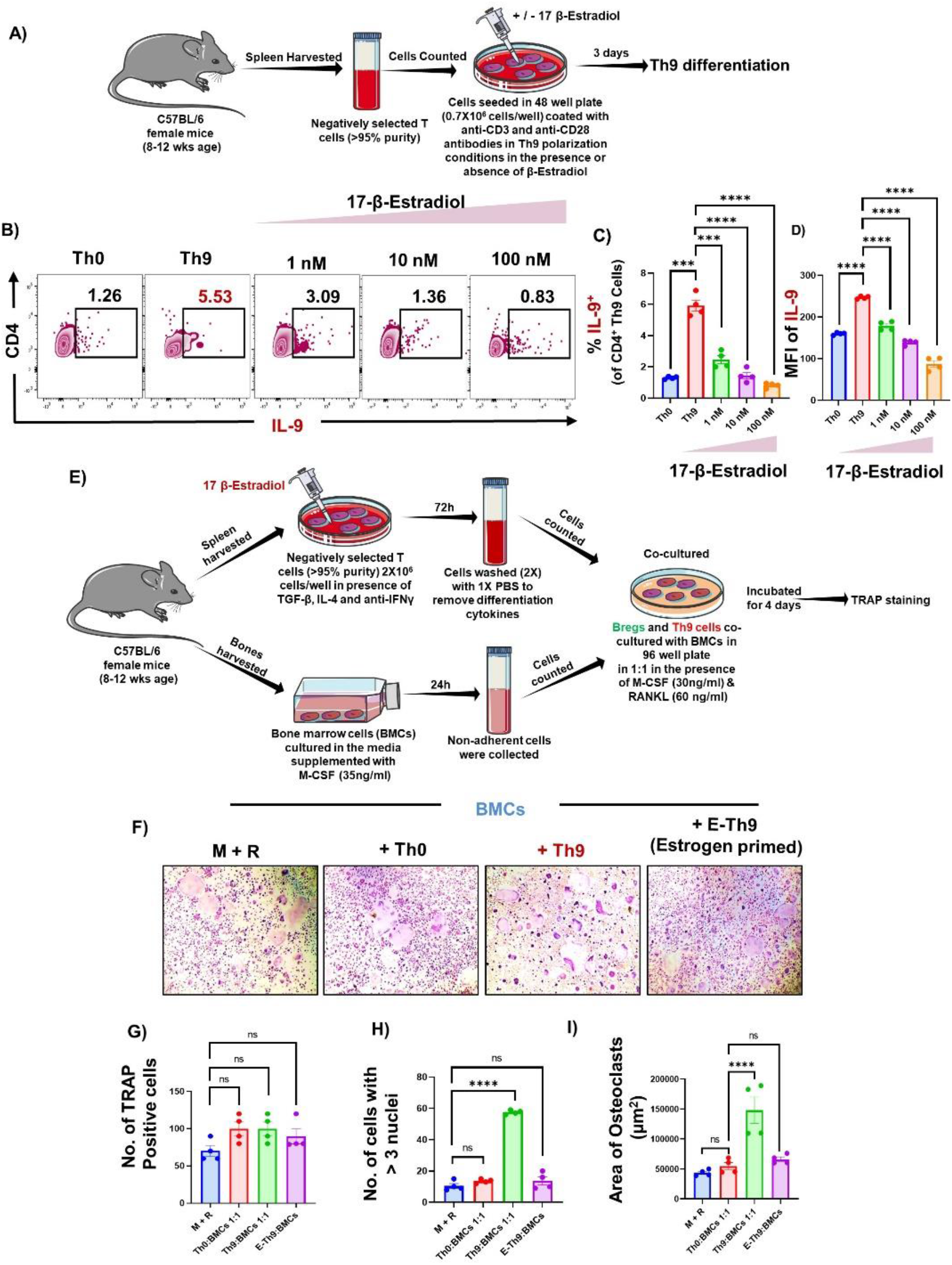
Th9 cells enhance osteoclastogenesis in an IL-9 dependent manner: A) Spleen was harvested and processed for negative selection of CD4^+^CD25^-^ naïve T cells. After estimation of CD4^+^ T cells purity (>95 %), naïve T cells were cultured in Th9 differentiation conditions for 72 hr. B) Zebra plot showing the differentiation of Th9 cells in the presence of 17-β Estradiol. C) Bar graphs representing the percentage of CD4^+^IL-9^+^ Th9 cells. D) MFI of IL-9. E) BMCs and Th0/Th9 were co-cultured in cell culture plate in the presence of M-CSF (30 ng/ml) and RANKL (60 ng/ml) for 4 days. Th9 cells were differentiated from naïve T cells for 72h prior to co-cultures. F) Photomicrographs represent TRAP positive cells. G) Number of TRAP positive cells. H) Number of TRAP positive cells with more than 3 nuclei. I) Area of osteoclasts. Data are expressed as mean ± SEM. Data were analyzed by unpaired student t test and analyzed by one way ANOVA. *P ≤ 0.05, **P ≤ 0.01, ***P ≤ 0.001, ****p ≤ 0.0001) compared with the indicated group.

### Th9 cells enhance osteoclastogenesis in an IL-9 dependent manner

We next investigated the direct role of Th9 cells on osteoclastogenesis by co-culturing BM cells with Th9 cells. Both the number and area of multinucleated osteoclasts (with > 3 nuclei) were significantly enhanced (5-fold) in Th9 cells co-cultured with BMCs in comparison to controls **(Fig. 5E-F)**, which was significantly reversed by E2 treatment **(Fig. 5F-I)**. We next sought to determine the mechanistic basis of osteoclastogenic Th9 cells. To ascertain this, we next examined the effect of IL-9 (signature cytokine of Th9 cells) on osteoclastogenesis (TRAP) *ex vivo*. IL-9 in a dose-dependent manner significantly enhanced osteoclastogenesis - higher number and area of TRAP-positive cells **(Fig. 6A-D)**. These data establish Th9 cells as an independent osteoclastogenic Th subset. Moreover, via stimulating osteoclastogenesis, Th9/IL-9 disrupts the balance of bone remodelling in favour of bone resorption.

**Fig 6:**
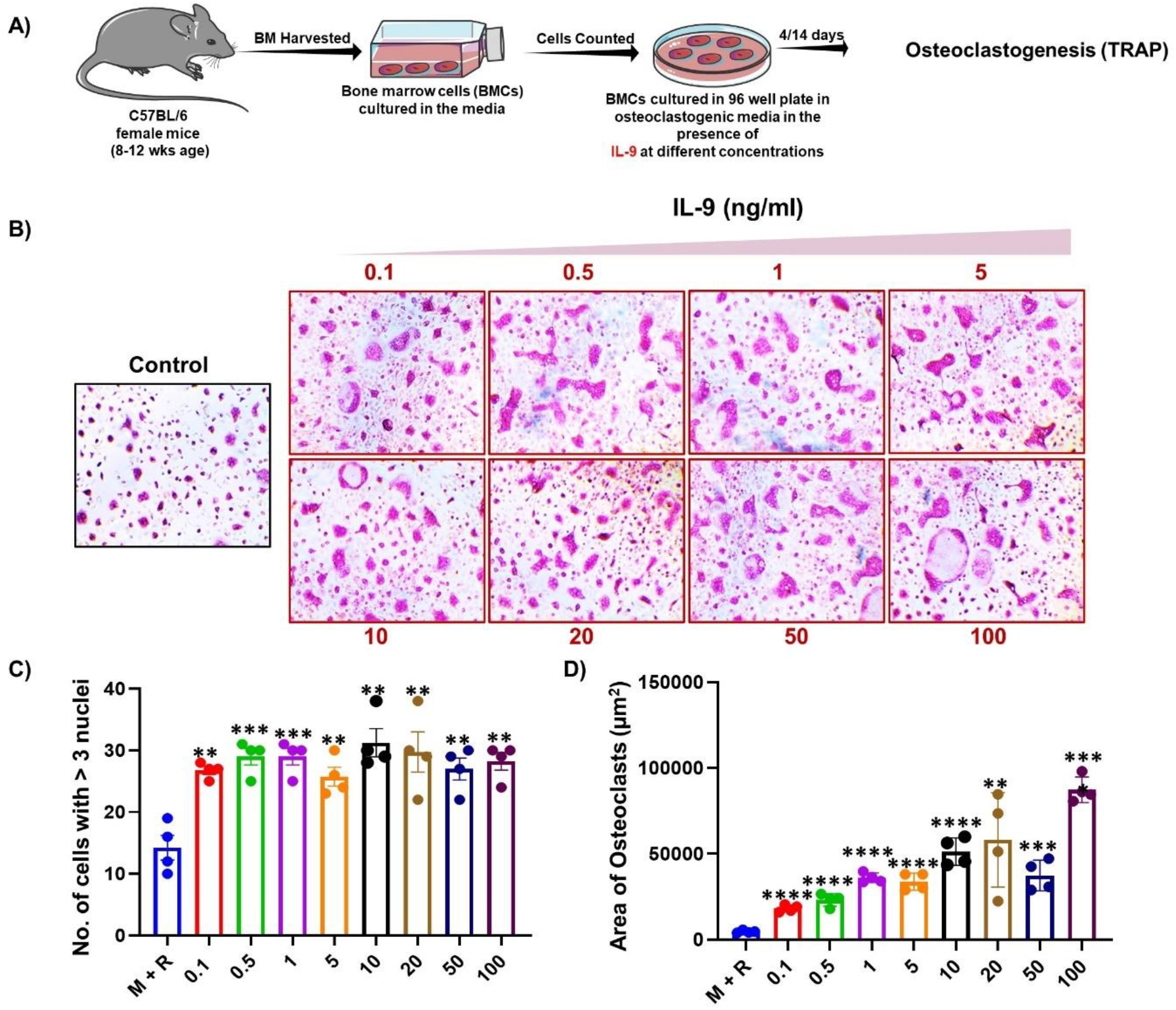
IL-9 enhances osteoclastogenesis: A) Osteoclast differentiation was induced in Bone Marrow cells (BMCs) with M-CSF (30 ng/ml) and RANKL (60 ng/ml) with or without IL-9 at different concentrations for 4 days. Giant multinucleated cells were stained with TRAP and cells with ≥ 3 nuclei were considered as mature osteoclasts. B) Photomicrographs at 20X magnification were taken. C) Number of TRAP positive cells with more than 3 nuclei. D) Area of osteoclasts. Data are expressed as mean ± SEM. Data were analyzed by unpaired student t test and analyzed by one way ANOVA. *P ≤ 0.05, **P ≤ 0.01, ***P ≤ 0.001, ****p ≤ 0.0001) compared with the indicated group.

### Anti-IL-9 therapy abrogates inflammatory bone loss under post-menopausal osteoporotic conditions

To conclusively demonstrate that IL-9 is involved in Ovx-induced osteoclastogenesis and consequent bone loss, we used anti-IL-9 antibody to neutralize this cytokine. Whereas isotype control failed to prevent bone loss in Ovx mice, anti-IL-9 antibody effectively protected against Ovx-induced bone loss. Ovx mice receiving anti-IL-9 antibody displayed significant decrease in osteoclastogenesis *ex vivo*. **(Fig. 7A-G)**, and mRNA levels of cathepsin K, the collagenase synthesized by mature osteoclasts for degrading collagen **(Fig. 7H)**. Moreover, the osteoclastogenic rheostat represented by *RANKL*:*OPG* ratio was significantly reduced upon anti-IL-9 antibody blockade **(Fig. 7I-K)**.

**Fig 7:**
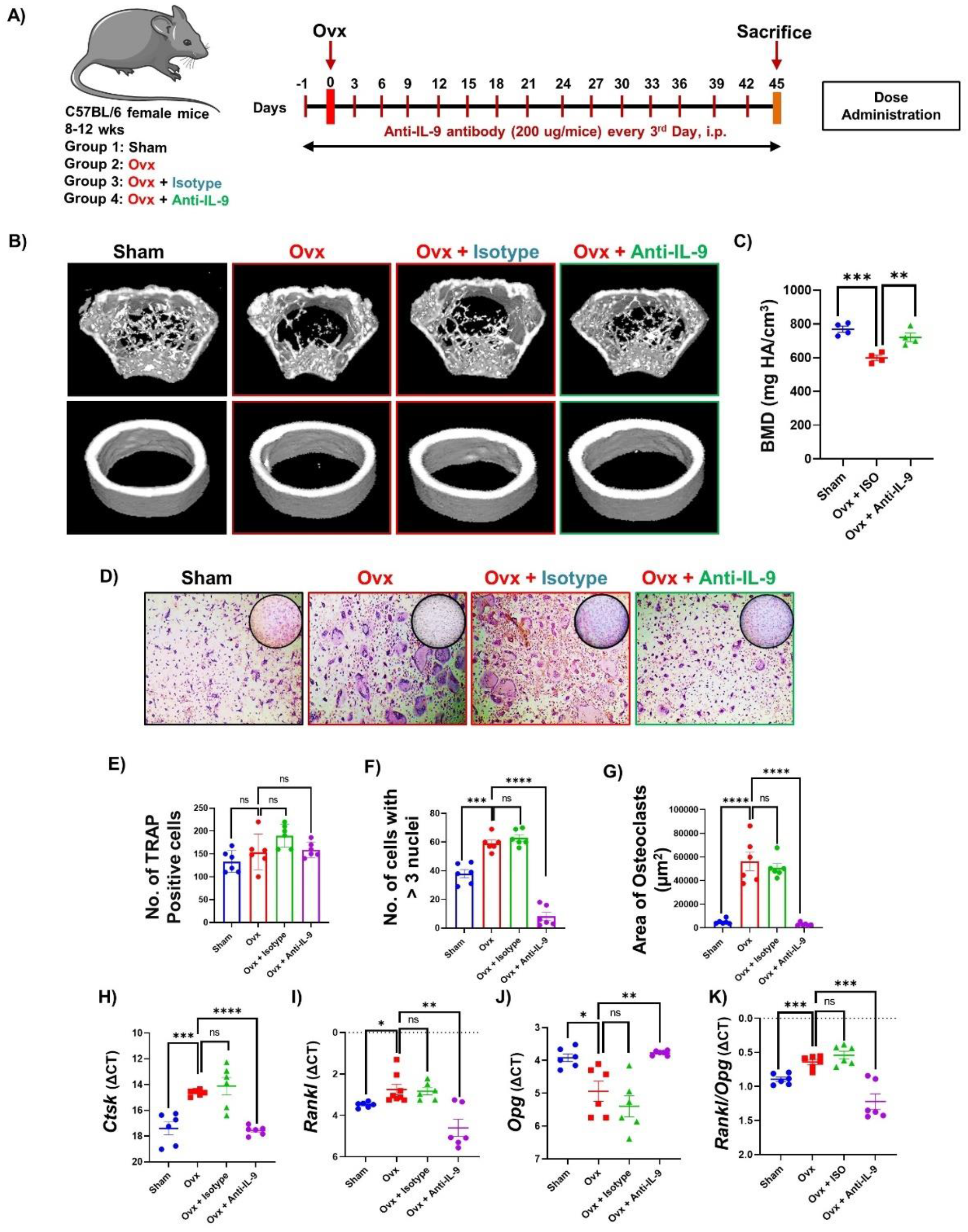
Anti-IL-9 therapy abrogates inflammatory bone loss under post-menopausal osteoporotic conditions: A) Experimental plan followed. B) Photomicrographs at 20X magnification were taken. C) Number of TRAP positive cells D) Number of TRAP positive osteoclasts with more than 3 nuclei. E) Area of osteoclasts. F) 3D micro-CT reconstructions of Femur metaphysis and femoral diaphyseal region. G) Bar graph representing relative expression of cathepsin K gene. H) Bar graph representing relative expression of RANKL. I) Bar graph representing relative expression of OPG gene. J) Bar graph representing RANKL/OPG ratio. Data are expressed as mean ± SEM. Data were analyzed by unpaired student t test and analyzed by one way ANOVA. *P ≤ 0.05, **P ≤ 0.01, ***P ≤ 0.001, ****p ≤ 0.0001) compared with the indicated group.

Since IL-9 induce the differentiation of Th17 cells **(Fig. 3F-H)**, we investigated the effect of blocking IL-9 on Th9 and Th17 cells in Ovx mice. Flow cytometric data revealed that anti-IL-9 antibody significantly reduced the percentage of both IL-9^+^ IL-17^-^ Th9 cells and IL-9^-^ IL-17^+^ Th17 cells in the BM of Ovx mice **(Fig. 8A-D)**, bringing their population below the sham levels **(Fig. 8A-D)**. Similar results were obtained in both spleen and peripheral blood of Ovx mice post-IL-9 blockade **(Fig. 8E-J)**. Consistent with these findings, the expression of IL-17A and Rorγt (Th17 transcription factor), along with FOXO-1 and IRF4 were significantly decreased upon IL-9 blockade in the BM of Ovx mice **(Fig. 8K-N)**. Together these data reveal that IL-9 blockade attenuates the differentiation/expansion of osteoclastogenic Th17 cells in Ovx mice. In animal models of autoimmune disorders, it has been demonstrated that IL-9 not only increases Th17 cell differentiation but also facilitates Th17 cell homing by elevating CCL20 expression. Our real-time results also showed that IL-9 inhibition markedly reduced *Ccl20* expression in BM, reducing osteoclastogenic Th17 cell homing **(Fig. 8O)**. Together, these findings show that, in post-menopausal osteoporotic conditions, IL-9 blockade preferentially inhibits differentiation/expansion and reduces the homing of osteoclastogenic Th17 cells, which further ameliorates the observed inflammatory bone loss.

**Fig 8:**
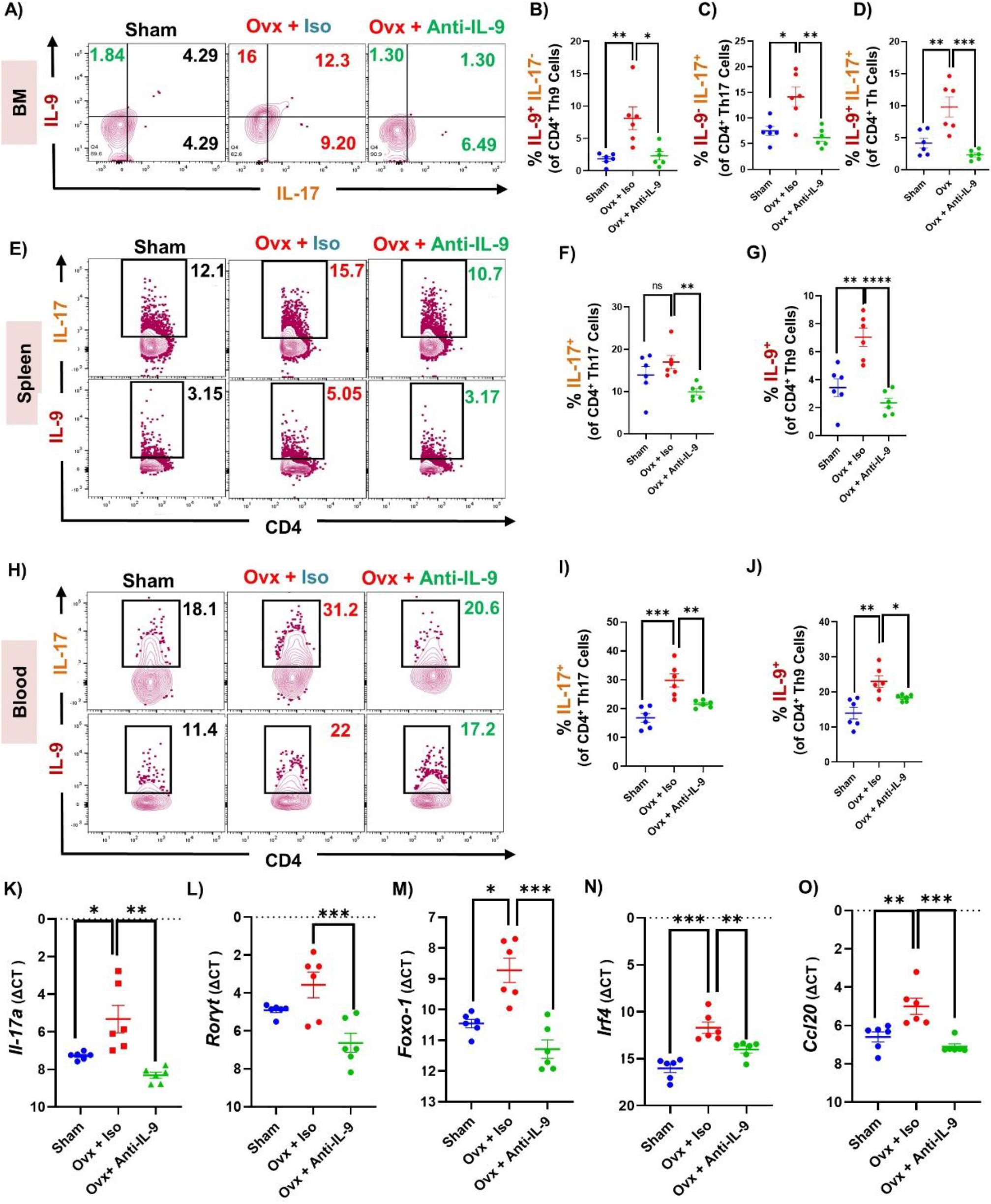
Blockade of IL-9 abrogates bone loss via modulating Th9 and Th17 cells: A) Contour plots representing CD4^+^IL-9^+^IL-17^+^ Th17 cells, CD4^+^IL-9^+^IL-17^-^ Th9 cells, CD4^+^IL-9^-^IL-17^+^ Th17 cells in BM. B) Individual plot representing CD4^+^IL-9^+^IL-17^-^ Th9 cells in BM. C) Individual plot representing CD4^+^IL-9^-^IL-17^+^ Th17 cells in BM. D) Individual plot representing CD4^+^IL-9^+^IL-17^+^ Th17 cells. E) Contour plot showing percentage of Th9 and Th17 cells in spleen. F) Percentage of Th17 cells in spleen. G) Percentage of Th9 cells in spleen. H) Contour plot showing percentage of Th9 and Th17 cells in blood. I) Percentage of Th17 cells in blood. J) Percentage of Th9 cells in blood. K) IL-17a gene expression. L) Rorγt gene expression. M) Foxo-1 gene expression. N) Interferon regulatory factor 4 (IRF4) gene expression. O) *Ccl 20* gene expression in BM. Data are expressed as mean ± SEM. Data were analyzed by unpaired student t test and analyzed by one way ANOVA. *P ≤ 0.05, **P ≤ 0.01, ***P ≤ 0.001, ****p ≤ 0.0001) compared with the indicated group.

### Both IL-9^+^ Th9 and IL-17^+^ Th17 cells are elevated in post-menopausal osteoporotic patients

We next sought to verify the pre-clinical results in postmenopausal osteoporotic patients. The clinical characteristics of the human subjects involved in the study are mentioned in the methodology section. Based on T scores, human subjects were categorized into either healthy control (HC, n = 10) and post-menopausal osteoporotic subjects (PMO, n = 10). There were significant increases in the levels of bone turnover markers, CTX-1 and osteocalcin in the osteoporotic group compared with HC group **(Fig 9A-B)**. Flow cytometry data revealed a significant enhancement in the percentage of both IL-9^+^ Th9 cells and IL-17^+^ Th17 cells in the PBMCs of PMO compared with HC **(Fig. 9C-I)**. Additionally, there was significant increase in the expression (MFI) of IL-9 and IL-17 in Th cells in the PMO compared with HC group. Embedded tSNE plot immunoprofiling further led to the identification of a unique cluster of IL-9 expressing Th cells in PMO subjects in comparison to HC **(Fig. 9C)**.

**Fig 9:**
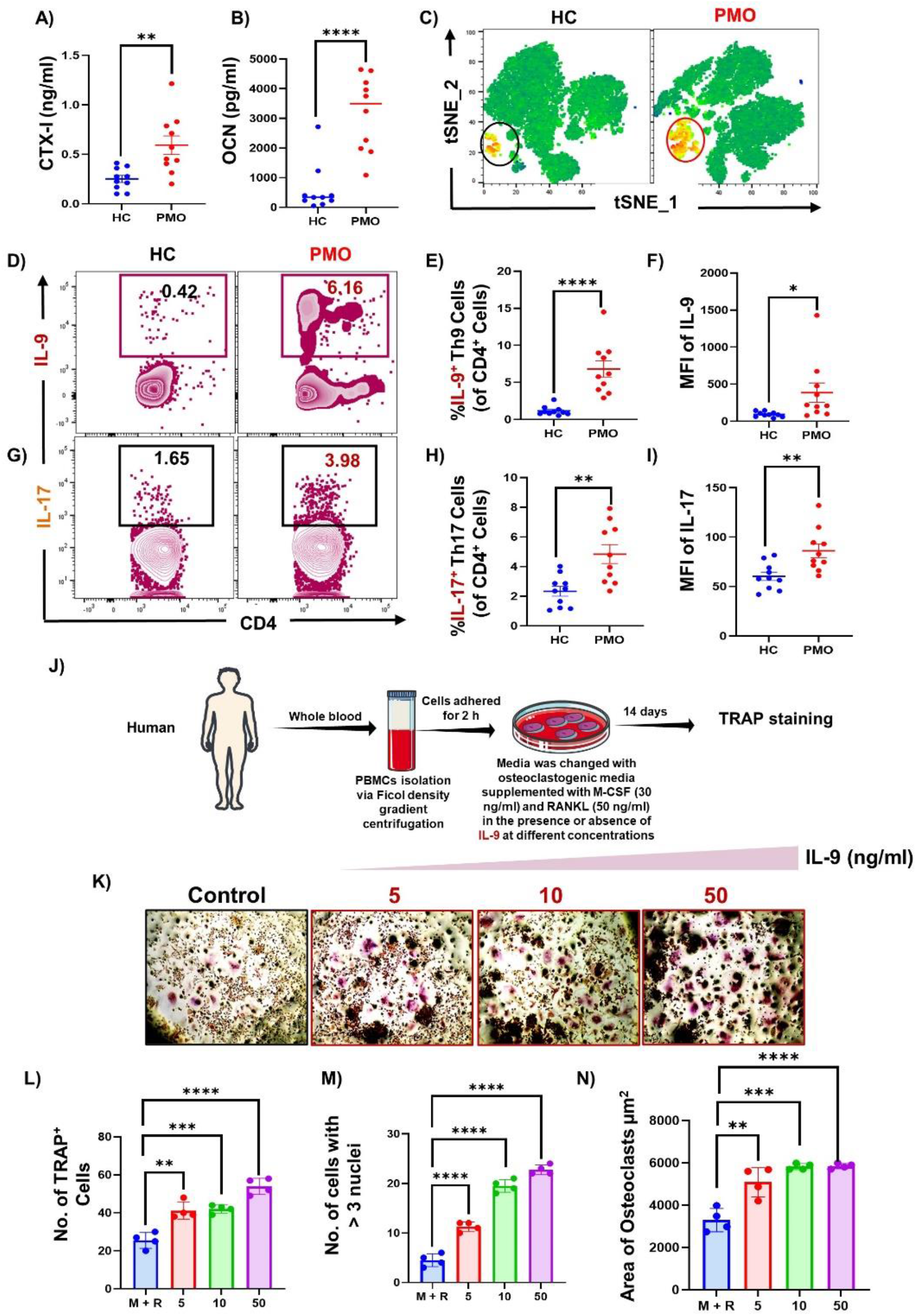
Both IL-9^+^ Th9 and IL-17^+^ Th17 cells are elevated in post-menopausal osteoporotic patients: A-B) ELISA data representing the levels of bone biochemical markers. C) tSNE plot for the PBMCs data set. D) Zebra plot representing the percentage of Th9 cells. E) Scatter plot representing the percentage of Th9 cells in control and PMO patients. F) MFI of IL-9 in CD4^+^ T cells in HC and PMO patients. G) Contour plot representing the percentage of Th17 cells in control and PMO patients. H) Scatter plot are representing percentage of Th17 cells. I) MFI of IL-17 in CD4^+^ T cells in HC and PMO patients. J) Osteoclast differentiation was induced in human PBMCs with M-CSF (25 ng/ml) and RANKL (100 ng/ml) with or without IL-9 cytokine at various concentrations for 14 days. Giant multinucleated cells were stained with TRAP and cells with ≥ 3 nuclei were considered as mature osteoclasts. K) Photomicrographs at 20 X magnifications were taken. L) Number of TRAP positive cells. M) Number of TRAP positive cells with more than 3 nuclei. N) Area of osteoclasts. Data are expressed as mean ± SEM. Data were analyzed by unpaired student t test and analyzed by one way ANOVA. *P ≤ 0.05, **P ≤ 0.01, ***P ≤ 0.001, ****p ≤ 0.0001) compared with the indicated group.

Moreover, consistent with the mice data, IL-9 also enhanced RANKL-mediated osteoclastogenesis in human PBMCs dose-dependently assessed by the number and area of TRAP positive multinucleated osteoclasts **(Fig. 9K-N)**. Altogether, results from the mouse and human studies established IL-9 as a key regulator of bone loss in PMO. These results identify IL-9 as a immune-therapeutic target for PMO.

## Discussion

The role of immune system in the pathophysiology of Osteoporosis i.e., “Immunoporosis” is now well established ^10,21,23^. Cytokines produced by Th cells play a vital role in modulating bone health under estrogen deficient conditions ^24–28^. Within the existing proinflammatory condition in Ovx mice marked by increase in inflammatory cytokines (IL-6 and TNF-α) and concurrent decrease in anti-inflammatory cytokines (IL-4, IL-10, and IFN-γ), rise in IL-9 further worsen the inflammatory burden. The simultaneous increase in IL-9, its receptor IL-9R, and Foxo-1 that promotes the differentiation of naïve CD4^+^ T cells into Th9 cells suggests the possibility of a positive feedback loop within bone marrow cells that could potentially amplify Th9 cell differentiation, creating an auto-reinforcement. Moreover, stimulation of Th17 differentiation by IL-9 adds another layer to the mechanisms involved in the proinflammatory state seen in OVX mice.

Numerous studies have shown that a lack of estrogen hormone causes bone loss by controlling a vast array of biochemical and immunological processes ^4,29,30^. The involvement of osteoclastogenic cytokines, such as IL-6, IL-17, and TNF-α, and anti-osteoclastogenic cytokines, such IL-4, IL-10, and interferon gamma (IFN-γ), in modulating bone health is noteworthy ^9,10,29,31^. Few studies have shown that elevated levels of the cytokine IL-9 in the serum of RA patients promote the growth of osteoclasts, raising the idea that blocking IL-9 might be a new immunotherapeutic target for preserving bone health in RA patients ^20,32^. In agreement with this, our results demonstrated that IL-9 cytokine levels were consistently elevated in Ovx mice. Further evidence that increased IL-9 levels may be a pertinent factor in inflammatory bone loss in estrogen-deficient situations comes from the fact that elevated IL-9 levels were observed to be inversely linked with BMD. Monocytes and macrophages, known progenitors of osteoclasts express the greatest levels of IL-9R ^33^. We also found that IL-9R expression was elevated in the BM (the primary site of osteoclastogenesis), which promoted osteoclastogenesis and resulted in IL-9-mediated inflammatory bone loss in Ovx mice model.

Age-related skeletal involution is indicated by decreased bone formation and increased BM obesity. Increased oxidative stress and decreased growth factor production, which activate transcription factors from the FOXO family, are associated with these changes. A study found that mice lacking FOXO1, 3, and 4 had more osteoblast progenitors than normal mice, which may indicate that FOXOs activation is a significant pathogenetic mechanism in the development of involutional osteoporosis ^34^. The ablation of FOXO-1 reduced the expression of IL-9 in both mouse and human Th9 and Th17 cells, demonstrating that FOXO-1 is a key transcription factor that is linked to the production of IL-9 by T helper cells in allergic asthma ^16,17,35,36^. In accordance with these results, we examined *Foxo-1* expression in the BM and found that it was significantly higher in Ovx mice. This shows that the observed increase in IL-9 levels in estrogen-deficient osteoporotic situations may have involved both Th9 and Th17 cells. A renaissance in interest in IL-9 biology has resulted from the revelation that multiple immune cell subsets, including ILC2, Th2 cells, Th9 cells, Th17 cells, and Tregs, express IL-9 ^37^. Surprisingly, we found that under estrogen deficient osteoporotic conditions, both Th9 and osteoclastogenic Th17 cells are the major sources of IL-9 cytokine. This was demonstrated by the noticeably higher levels of IL-9^+^IL-17^-^ Th9 cells, IL-9^-^IL-17^+^ Th17 cells, and IL-9^+^IL-17^+^ T cells in the BM under estrogen-deficient circumstances.

Our results concur with studies showing that Th17 cells are the osteoclastogenic T cell subset that aggravates bone loss in estrogen-deficient conditions ^7,10,38–42^. Published studies reported that IL-9 along with TGF-β promotes the differentiation of naïve T cells towards Th17 phenotype ^43^. During the development process, Th9 cells produce the cytokine IL-9, which interacts with TGF-β to differentiate naive CD4^+^ T cells into Th17 cells. This starts an autocrine feedback loop, which further accelerates the proliferation of Th17 cells ^44^. The important role of the IL-9 cytokine in the differentiation of Th17 cells is highlighted by the considerable reduction in the development to Th17 cells in IL-9R^-/-^ CD4^+^ T cell when compared to wild type ^43,44^. In an *in vivo* temporal kinetics analysis, we found that estrogen deprivation triggers the release of IL-9 by Th cells, which further boosts the proportion of Th17 cells that express IL-17 in Ovx mice. Similar findings were made in our *ex vivo* study, where we found that estrogen counteracts both the effect of IL-9 on Th17 cell differentiation as well as decreasing Th9’s capacity to produce IL-9 under Th9 differentiation conditions. These findings strongly imply that the presence of estrogen is sufficient under physiological circumstances to prevent the formation of Th9 cells, which further prevents the differentiation of Th17 cells. Th17 cells are an established osteoclastogenic subset of Th cells but the role of Th9 cells on osteoclastogenesis is still warranted ^39^. Th9 cells significantly boost osteoclast development *ex vivo* in an IL-9-dependent manner, thereby establishing themselves as the newest member of the growing family of osteoclastogenic Th cells. Notably, we further discovered that estrogen primed Th9 cells were incapable of inducing osteoclastogenesis *ex vivo*.

According to studies, estrogen deprivation causes Th17 cells to migrate to the BM in a CCR6-CCL20 dependent way ^45,46^. These ligands are prominently expressed by BM stromal cells and hemopoietic cells as well as at various inflammatory sites. The chemokine ligand CCL20, which binds to CCR6, is responsible for mediating the influx of Th17 cells expressing CCR6 into the BM ^47,48^. Additionally, research in a mouse model of experimental autoimmune encephalomyelitis (EAE) revealed that IL-9 plays a role in the homing of Th17 cells to the central nervous system ^49^. These findings thus raise the notion that increased IL-9 levels brought on by estrogen deprivation may be one of the main drivers of osteoclastogenic Th17 cell homing to the BM. Surprisingly, we discovered that inhibiting IL-9 causes a significant reduction in CCL20 expression in the BM of Ovx mice, which was further associated with a decline in the number of Th17 cells in the BM. Unexpectedly, we also observed a substantial decline in the number of Th9 cells in the BM of Ovx mice following IL-9 suppression. In addition, both IRF4 and FOXO-1 were found to be significantly reduced in the Ovx mice post IL-9 blockade, thereby attenuating the expansion of Th9 cells. Our findings imply that IL-9-producing Th cells contribute negatively to inflammatory bone loss and that functional IL-9 blockade may be used therapeutically to enhance bone health in estrogen-deficient environments. A major conclusion of the study was that anti-IL-9 therapy enhanced bone health by preserving bone microarchitecture, controlling osteoclast development, and regulating the levels of bone biochemical markers linked to the bone remodelling process, including TRAP, RANKL, OPG, and cathepsin-K. According to our study, IL-9 is necessary for the development of Th17 cells. These findings suggested that blocking IL-9 would also have an impact on Th17 cells, hence reducing the associated bone loss reported in PMO.

Th9 cells stimulated the formation or activation of osteoclasts in a co-culture system, suggesting its role in promoting osteoclast activity and bone loss. Estrogen blocks Th9-induced Th17 differentiation and Th9 differentiation to produce IL-9 thus limiting the subsequent promotion of osteoclast activity in Ovx mice. Our findings in the mouse model are supported by similar observations made in humans where IL-9^+^ Th9 cells and IL-17^+^ Th17 cells in the PBMCs of PMO were significantly higher compared with control. Moreover, IL-9 and IL-17 expression were enhanced in Th cells and presence of a unique cluster of Th cells expressing IL-9 in individuals with osteoporosis compared to a control group. At the functional level, IL-9 exacerbated the RANKL-induced differentiation of human PBMCs to osteoclasts thus highlighting its mediatory role in osteoclastogenesis in humans. IL-9 blockade by antibody attenuates bone loss in Ovx mice with attendant decrease in IL-9, IL-17 and CCL-20. These cytokines may induce osteoblasts to produce RANKL, a key cytokine that promotes osteoclast differentiation and anti-IL-9 antibody downregulated the Ovx-induced rise in RANKL:OPG ratio.

Targeting IL-9, given its position upstream in the inflammatory cascade including IL-17, CCL-20 and RANKL that leads to bone loss, represents an appealing and potentially superior strategy to existing anti-resorptive therapies, particularly denosumab, the neutralizing antibody against RANKL that turns off the bone remodelling cycle. As a result, denosumab suffers from rapid bone turnover rebound after discontinuation leading to increased risk of multiple vertebral fractures, especially in patients who have other risk factors for fracture. By neutralizing IL-9 it is possible to specifically modulate the intensity and duration of inflammation without inducing toxicity of various vital organs **(Fig, S5A)**.

In the current study, we identified the mechanisms by which IL-9 could influence bone health in estrogen-deficient environments. It is still unclear if Th9, in the absence of Th17, can cause bone loss in a pre-clinical osteoporosis mouse model. Future studies could also examine the effects of inhibiting the chemotactic potential of CCL20 by reducing Th17’s homing to the BM. Undoubtedly, our findings demonstrated the presence of IL-9^+^IL-17^+^ Th cells in the BM of Ovx mice, but a fascinating area for future study would be the precise features and function of this population in PMO.

This fine-tuning allows for the reduction of inflammation associated with bone loss while preserving essential aspects of the immune response. In summary, this study offers the first evidence of the osteoclastogenic properties of Th9 cells, which are mediated by IL-9 (Fig. 10). Additionally, our study demonstrated that both inflammatory Th9 and Th17 cells sustain the low-grade chronic inflammation observed in osteoporotic situations. In conclusion, our pre-clinical and clinical data substantiate Th9 cells’ osteoclastogenic and osteoporotic functions in estrogen-deficient conditions and add Th9 to the growing list of osteoclastogenic T helper subsets. These results highlight the revolutionary relevance of IL-9/Th9 as a promising immunotherapeutic target for the management of inflammatory bone loss disorders observed in osteoporosis.

**Fig 10:**
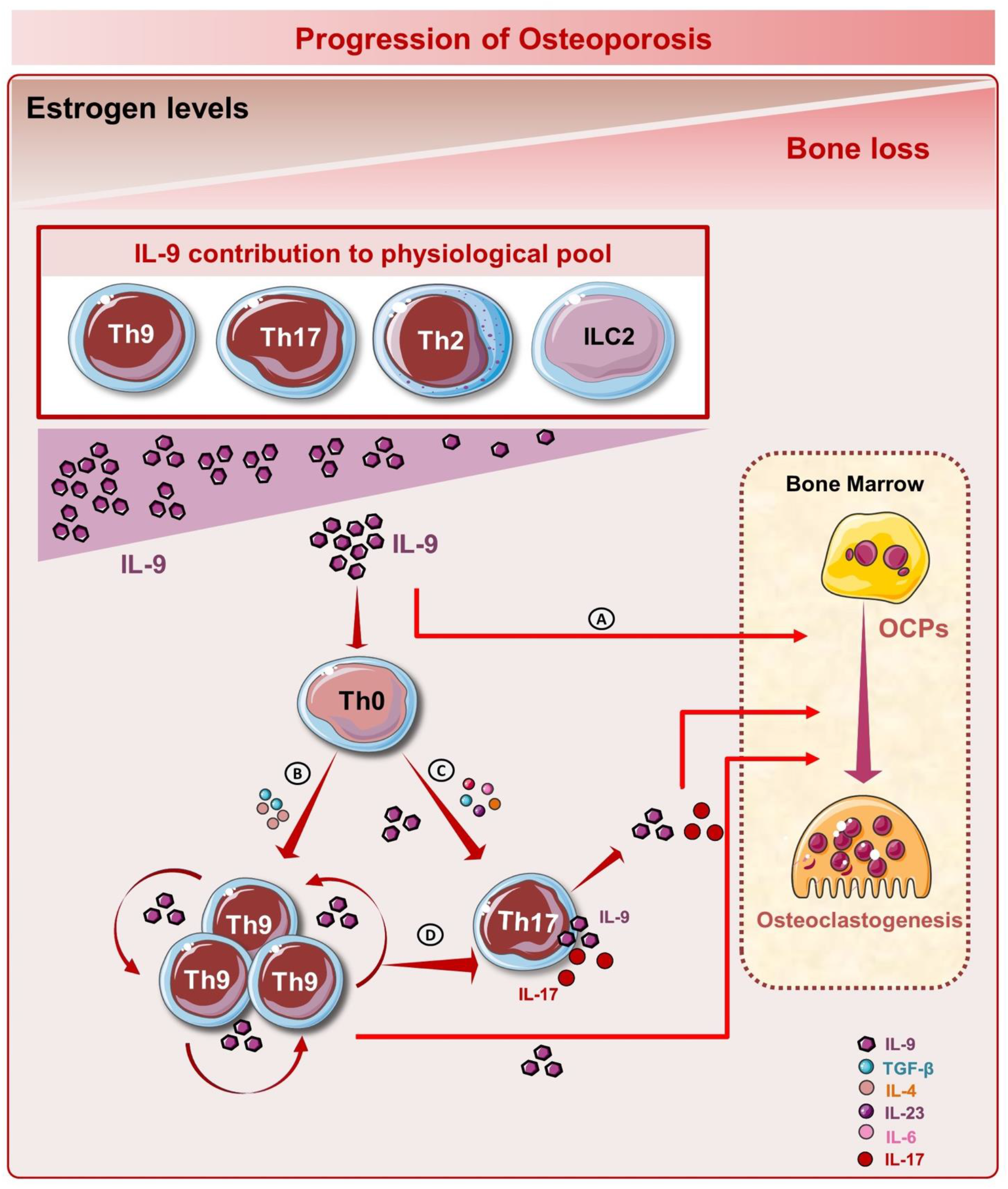
Summary of results: Among other innate and adaptive immune cells Th9 and Th17 cells are predominant source of IL-9 in Ovx mice. A) IL-9 induces the differentiation of BMCs to mature osteoclasts. B-D) IL-9 induces the expansion of Th9 cells and differentiation of Th17 cells that further via inducing secretion of IL-9 and Th17 cells induces the differentiation of osteoclasts. These findings suggest that anti-IL-9 therapy improves bone health in a preclinical model of postmenopausal osteoporosis by limiting the development of osteoclasts and osteoclastogenic Th9/Th17 cells.

## Material and Methods

### Reagents and Antibodies

The following antibodies were procured from Biolegend (USA): PerCp-Cy5.5-Anti-Mouse-CD4-(RM4-5) (550954), BV786-Anti-Mouse-IL-17A-(506928), BV421-Anti-Mouse-IL-9-(514109), BV711-Anti-Mouse-IL-4-(504133), BV605-Anti-Mouse-IFNγ-(505839) etc. Foxp3/Transcription factor staining buffer (0-5523-00) and RBC lysis buffer (00-4300-54) were purchased from eBioscience. Acid phosphatase, leukocyte (TRAP) kit (387A) was purchased from Sigma (USA). Human TGF-β1 (AF-100-21C), IL-9 (219-19) and IL-4 were procured from PeproTech (USA). Anti-CD3 and Anti-CD28 were purchased from BD (USA). α-Minimal essential media (MEM) and RPMI-1640 were purchased from Gibco (USA). Following ELISA kits were brought from R&D: Mouse IL-10 (M1000B) and Mouse IL-17 (M1700) Quantikine ELISA kits. The Following ELISA kits and reagents were brought from BD (USA): Mouse IL-6 (OptEIA^TM^-555240), Mouse TNF-α (OptEIA^TM^-560478) and Mouse IL-9. Anti-IL-9 was purchased from BioXcell (USA).

### Post-menopausal osteoporotic mice model

All in vitro and in vivo studies were performed on female C57BL/6 mice (8 to 10 weeks old). All the mice were kept in specified pathogen-free (SPF) environments at the animal facility of All India Institute of Medical Sciences (AIIMS), New-Delhi-India. For the investigation and kinetic study, following groups: sham (control-ovaries intact) and ovariectomized (ovx) (n=6/grp) were employed for 3 time periods (15, 30 and 45 days). A surgery stimulus procedure was performed on a healthy control (sham) group. Mice in the ovx group underwent bilateral-ovariectomy after anesthetizing them with ketamine (100-150 mg/kg) and xylazine (5 mg/kg) intraperitoneally. Both groups were kept in polycarbonate cages on a 12-hour light/dark cycle and were given access to sterile food and autoclaved water as needed. After respective time periods, mice were euthanized by CO_2_ asphyxiation and blood, bones and lymphoid tissues especially bone marrow (BM), spleen and mesenteric lymph nodes (MLN) were harvested for further analysis. For IL-9 blockade, mice were injected intra peritoneally (i.p) with 200 μg/mice every other day starting on day -1 of Ovx surgery for 45 days. Another group received isotype-matched IgG at a dose of 200 ug/ mice on day -1 of Ovx surgery for day 45. All procedures were carried out in accordance with the guiding principles, suggestions, and after the protocol was approved by the institutional animal ethics committee of AIIMS, New Delhi, India (384/IAEC-1/2022).

### Scanning Electron Microscopy (SEM)

As previously reported ^21,22^ SEM was done for the cortical area of the femoral bones. Briefly, bone samples were stored in 1 % Triton-X-100 for 2 to 3 days before being transferred to 1 % PBS buffer till final analysis. After forming the bone slices, the samples were dried under an incandescent lamp before being coated with sputter. Then, using a Leo 435 VP microscope with a 35-mm photographic system, bones were scanned. To capture the tiniest cortical region, SEM pictures were digitally captured at a 100 X magnification. Following imaging, MATLAB was used to further analyse the SEM pictures (MathWorks, Natick, MA, USA).

### Micro-computed tomography (µ-CT)

Micro CT (µ-CT) scanning, and analysis were done by employing in vivo X-ray SKYSCAN 1076 scanner (Aartselaar, Belgium) tomography. First, samples were positioned correctly in the sample holder before scanning was conducted at 50 kV, 204 mA, and a 0.5 mm aluminium filter. NRecon software was used for the reconstruction procedure. Following reconstruction, 100 slices of the secondary spongiosa at 1.5 mm from the distal edge of the growth plates were used to draw the ROI. Bone sample micro-architectural parameters are evaluated and computed using CTA software. The measurements of bone volume/tissue volume (BV/TV), trabecular thickness (Tb. Th), trabecular separation (Tb. Sp), and other 3D-histomorphometric parameters were obtained. The bone mineral density (BMD) of the LV5, femur, and tibia were calculated using the volume of interest from u-CT scans made for trabecular and cortical regions. Using calibrator hydroxyapatite phantom rods with known BMD (0.25 g/cm^3^ and 0.75 g/cm^3^), BMD was calculated ^21^.

### Histological analysis

For decalcification, right femoral bone was extracted and fixed in 4 % paraformaldehyde (PFA) for 24 h. After fixation, bone was decalcified by placing the bone in 10 % tetrasodium EDTA aqueous solution on a rocker for 15 days. Paraffin-embedded sections (5 μm) from femur were processed for the histological haematoxylin (H) and eosin (E) staining analysis. Sections were imaged using a microscope.

### Th9 and Th17 cell differentiation assay

Splenic naïve T cells from C57BL/6 mice were purified by magnetic separation described previously ^21^. Briefly, after RBC lysis cells were subjected to a T cell enrichment cocktail (BD, USA) and untouched negatively selected CD4^+^CD25^-^ naïve T cells (purity > 95 %) were seeded in anti-CD3 (10 µg/ml) and anti-CD28 (2 µg/ml) mAbs coated 48 well plate. For Th9 cell differentiation, cells were cultured in RPMI-1640 media and polarized with IL-4 (20 ng/ml), TGF-β1 (5 ng/ml) and anti-IFN-γ (5 µg/ml) in the presence or absence of 17β-estradiol (1nM, 10 nM and 100 nM) and incubated for 3 days. For Th17 cell differentiation, naïve T cells were stimulated with TGF-β1 (2 ng/ml), IL-6 (30 ng/ml), IL-23 (20 ng/ml), anti-IL-4 (10 µg/ml), and anti-IFN-γ (10 µg/ml) and incubated for 3 days.

At the end of incubation, cells were harvested, and flow cytometry was performed for estimating the percentages of CD4^+^IL-9^+^ Th9 cells. In addition, cells were harvested and washed thrice with 1X PBS to remove the traces of differentiating cytokines before co-culturing with the BMCs for osteoclast differentiation assay.

### Co-culturing of Th9 cells with BMCs for osteoclastogenesis

As previously mentioned, mouse BMCs were used to generate osteoclasts. In a nutshell, BMCs were harvested from the femur and tibiae of 8–10-week-old mice, and RBC lysis was carried out by using 1X RBC lysis buffer. Following RBC lysis, cells were cultured for overnight in a T-25 flask using endotoxin-free-MEM media (10 % FBS) supplemented with M-CSF (35 ng/ml). Next day, non-adherent cells (BMCs) were collected and co-cultured with Th9 cells or Th0 cells in a 96 well plate at 1:1 cell ratio for 4 days in the presence of M-CSF (30 ng/ml) and RANKL (60 ng/ml). The media was replenished every two days with new media supplemented with fresh M-CSF and RANKL factors. The tartrate resistant acid phosphatase (TRAP) staining procedure was carried out after 4 days of incubation.

### Osteoclasts differentiation and tartrate resistant acid phosphatase (TRAP) staining

For osteoclastogenesis, mouse BMCs were harvested from 8- to 12-week-old C57BL/6J mice by flushing the femoral bone with α-MEM media. Following RBC lysis with 1 X RBC lysis buffer, BMCs were cultured overnight in a T-25 flask in endotoxin-free complete α-MEM media (10 % heat-inactivated FBS) with M-CSF supplemented at 35 ng/ml concentration. The next day, non-adherent cells were collected and seeded in 96-well plates (50,000 cells/well) in complete α-MEM media supplemented with osteoclastogenic factors including M-CSF (30 ng/ml) and RANKL (60 ng/ ml) in the presence or absence of IL-9 cytokine at different concentrations: 0.1 ng/ml, 0.5 ng/ml, 1 ng/ml, 5 ng/ml, 10 ng/ml, 20 ng/ml, 50 ng/ml and 100 ng/ml for 4 days. After 72 h, half media was replenished with fresh complete α-MEM media supplemented with fresh factors.

Lastly for assessing the generation of multinucleated osteoclasts, TRAP staining was carried out according to the manufacturer’s recommendations. After incubation, cells were briefly washed three times with 1XPBS and then fixed for 10 minutes at 37°C in a fixative solution made of citrate, acetone, and 3.7 % formaldehyde. Fixed cells were stained for TRAP at 37 °C in the dark for 5–15 minutes after being washed twice with 1XPBS. Osteoclasts that have ≥ 3 nuclei and TRAP positive were regarded as mature osteoclasts. Multinucleated TRAP-positive cells were also counted and seen using an inverted microscope (ECLIPSE, TS100, Nikon). The area of the TRAP-positive cells was measured using Image J software (NIH, USA).

### Flow cytometry

For intracellular cytokine analysis, cells were harvested from various lymphoid organs and suspended in complete RPMI-1640 medium (R10) were seeded at a density of 1 X 10^6^ cells/ml in 96 well plate. Following that, cells were stimulated for the following 5h using a combination of protein transport inhibitor (Monensin) (Biolegend, USA), Ionomycin (1 ug/ml), and PMA (50 ng/ml, Sigma Aldrich). After that, cells were harvested and stained for specific cell surface and intracellular markers. Flow cytometry analysis was performed according to the previously defined methodologies ^22^. For evaluating the percentage of Th9 and Th17 cells flow cytometry was employed. First, cells were surface stained for Th9 and Th17 cells (anti-CD4-PerCP-Cy5.5) and incubated for 30 minutes in the dark on ice. Cells were washed, fixed, and permeabilized with 1X fixation permeabilization buffer following surface staining. Finally, intracellular staining was performed, and cells were further stained with BV421-conjugated-anti-IL-9 and BV786-conjugated-anti-IL-17 antibody. After washing, cells were acquired on BD FACSymphony (USA). The samples were then examined using Flow Jo 10 (Tree Star, Woodburn, OR, USA) software.

### Study Subjects

After being screened using Dual Energy X-Ray Absorptiometry (DEXA), clinicians involved in the study have chosen the study participants. Women who had a history of smoking, drinking, endocrinopathy, diabetes, or hypertension, were pregnant or nursing, taking oral contraceptives, glucocorticoids, hormone replacement therapy, or were taking any medications that affected bone mass or the immune system were excluded from the study. Inclusion criteria for the study included normotensive, physically active, non-smoker, and non-drinker women. Prior to enrolment, postmenopausal women had experienced natural menopause for at least a year. Each participant provided their written, informed consent before having their clinical history documented. All the measures were carried out following the proper clearance of the protocols filed to the institute Ethics Committee for Post Graduate Research (IECPG-482), AIIMS, New Delhi, India. In accordance with WHO guidelines, the chosen postmenopausal women were categorised as normal (T score ≥ -1.0 at both sites), osteopenic (T score between -1.0 and -2.5 at any site), and osteoporotic (T score ≤ -2.5 at any site). Total 20 patients were recruited: 10 in each group in both healthy control and post-menopausal osteoporotic (PMO) subjects. Each participant gave consent to the drawing of 10 ml of blood into heparinized tubes. Prior to usage, aliquots of the plasma and serum samples were stored at -80 °C. PBMCs, isolated from the blood were used immediately for flow cytometry and transcriptional analysis.

### Quantitative PCR (q-PCR)

RNA was extracted from the lung tissue cells using the RNeasy Mini Kit (Qiagen, USA) according to the manufacturer’s instructions. The RNA quality was checked via 28s and 18s RNA using Bio Analyzer (Agilent Technologies, Singapore). Only samples with optimum RNA Integration Number value of ≥ 7 were used. RNA was also quantified using spectrophotometer (Nanodrop Technologies, USA). For complimentary DNA conversion, 1µg of total RNA was used. First Strand c-DNA synthesis kit (Thermo Scientific, USA) was used according to manufactures instructions. Further, gene expression was measured using quantitative real-time (Applied Biosystems, Quantstudio^TM^-5, USA). Triplicate samples of cDNA from each group were amplified with customized primers viz. IL-9, IL-9R, IL-17A, ROR-γt, IRF4, FOXO-1, Cathepsin K, RANKL, OPG and CCL20 normalized with arithmetic mean of GAPDH housekeeping gene. 25 ng of c-DNA was used per reaction in each well containing the 2X SYBR green PCR master mix (Promega, USA) along with appropriate primers. Threshold cycles values were normalized and expressed as relative gene expression.

### Human osteoclast differentiation from PBMCs

To obtain PBMCs, heparinized blood was gently layered on Histopaque at 1:3 dilution in a 15 ml tube and later centrifuged at RT for 25 min at 800 X g (brake off). Buffy coat layer underlying the plasma was carefully collected in fresh tube, washed with 1X PBS, gently inverted and centrifuged at 400 X g for 10 min at 4°C. The obtained PBMCs were seeded at a density of 1X 10^6^ cells per well in a 96-well plate, incubated for 2 hours in a humidified 5 % CO_2_ incubator and later washed twice with α-MEM (without FBS).

As soon as cells adhered, they were incubated with complete α-MEM media (10 % FBS) supplemented with M-CSF (30 ng/ml) and RANKL (100 ng/ml) in the presence or absence of IL-9 cytokine at various concentrations. The plate was then incubated for the following 14 days in a CO_2_ incubator with half media replenishment every third day (i.e., 72 h). Following incubation, multinucleated osteoclast development was assessed using TRAP staining.

### Statistical analysis

GraphPad Prism version 9.0 was used for statistical analysis (San Diego, California, USA). Data were represented as mean ± SEM and *p* ≤ 0.05 value was regarded as statistically significant. Unpaired parametric Student’s t test was used to compare the two groups. For in vitro experiments (TRAP and Th9 cell differentiation assay), we used 3 biological replicates and 3 technical replicates. For the statistical analysis of human patient sample data, Mann-Whitney test was carried out.

#### Study approval

All clinical investigation conducted according to Declaration of Helsinki principles. All human studies have been approved by the appropriate institutional review board and written informed consent was received prior to participation.

#### Data availability

Relevant information about data availability directly from the corresponding author.

## Author contributions

RKS: conceptualization, methodology, validation, formal analysis, resources, writing original draft, visualization, supervision, project administration and funding acquisition. LS: Data curation, writing original draft, investigation, and formal analysis. CS: Data curation and formal analysis. PKM: methodology and formal analysis. NC: methodology, writing-review and editing and formal analysis.

## Acknowledgment

This work was financially supported by the following projects: DBT(BT/PR41958/MED/97/524/2021), ICMR (61/05/2022-IMM/BMS) Govt. of India and AIIMS, Intramural project (AC-939) New Delhi-India sanctioned to RKS. LS, CS and RKS acknowledge the Department of Biotechnology AIIMS, New Delhi-India for providing infrastructural facilities. LS thank UGC thank for research fellowship.

## Notes

### Competing Interest Statement

The authors have declared no competing interest.

